# Biased phylodynamic inferences from analysing clusters of viral sequences

**DOI:** 10.1101/095661

**Authors:** Bethany L. Dearlove, Fei Xiang, Simon D. W. Frost

## Abstract

Phylogenetic methods are being increasingly used to help understand the transmission dynamics of measurably evolving viruses, including HIV. Clusters of highly similar sequences are often observed, which appear to follow a ‘power law’ behaviour, with a small number of very large clusters. These clusters may help to identify subpopulations in an epidemic, and inform where intervention strategies should be implemented. However, clustering of samples does not necessarily imply the presence of a subpopulation with high transmission rates, as groups of closely related viruses can also occur due to non-epidemiological effects such as over-sampling. It is important to ensure that observed phylogenetic clustering reflects true heterogeneity in the transmitting population, and is not being driven by non-epidemiological effects.

We quantify the effect of using a falsely identified ‘transmission cluster’ of sequences to estimate phylodynamic parameters including the effective population size and exponential growth rate. Our simulation studies show that taking the maximum size cluster to re-estimate parameters from trees simulated under a randomly mixing, constant population size coalescent process systematically underestimates the overall effective population size. In addition, the transmission cluster wrongly resembles an exponential or logistic growth model 95% of the time. We also illustrate the consequences of false clusters in exponentially growing coalescent and birth-death trees, where again, the growth rate is skewed upwards. This has clear implications for identifying clusters in large viral databases, where a false cluster could result in wasted intervention resources.

## Introduction

Boosted by the increasing use of viral genotyping for clinical purposes, there are more HIV sequence data than ever, representing around a third of all viral sequences in Genbank, which can also be used to characterise the transmission dynamics of HIV. Grouping sequences into phylogenetic clusters has previously proven useful for sifting through these large datasets, and this approach has been used to correlate transmission with contact rates, social network structures, risk behaviours and presence of co-infections with other viruses in a number of HIV studies, including those from the United Kingdom, Switzerland, Canada, the Netherlands and South America [1–5]. However, the definition of a phylogenetic cluster has not so far been standardised, despite there being numerous approaches and software implementations to identify them [6–10].

In standard epidemiology, a cluster is defined as a higher than expected burden or incidence of disease in close proximity in time and space [11]. In phylogenetics, a cluster is simply a group of closely related sequences, or a subtree within the full phylogeny, linked by a single recent common ancestor. This definition on its own does not help resolve epidemiological links, since broad clustering in a tree of all HIV-1 sequences will simply identify subtypes [10]. However, this does not mean to say that the phylogenetic and epidemiological definitions of a cluster cannot be reconciled; sequences that are more closely related genetically may have a smaller distance in terms of the number of transmission events that have occurred between them, all other things being equal. Thus the identification of phylogenetic clusters can represent groups of recent transmission when identified using a measure of relatedness. Whilst exact definitions vary in the literature, this relatedness, referred to here as the threshold, is usually in the form of an allowable distance between sequences, either through a cut-off of genetic diversity, or in a time period reflective of time to diagnosis. Methods using the mean, median and maximum distance for the threshold have been proposed [1,7,12]. Constraints on the certainty of the subtree (through a bootstrap or posterior probability, for example), subtree size, and geography of isolates have also been introduced [2,13].

Regardless of how clusters are identified, resulting clusters are undirected between pairs of sequences, and thus can also be drawn as a network (Figure 1b). The sizes of clusters can be represented as a (component) distribution (Figure 1c). The distribution of cluster sizes is often found to be right skewed, with a large number of very small clusters and fewer large ones, shown to be well-fitted by a power-law distribution [1,12]. This shape is often thought to be a reflection of the heterogeneity in transmission, particularly for sexually-transmitted infections where a similar pattern of skew can be seen in risky sexual contacts [14]. Cluster sizes which go against the power law behaviour, such as the cluster of size 79 in Figure 1, tend to be of particular interest since these suggest new or unexpected patterns of transmission for further analysis. However, without a null model in which there is no heterogeneity in transmission, it is difficult to interpret these distributions. Indeed, there is nothing special about the largest cluster in Figure 1, since the phylogeny was simulated using a randomly mixing constant coalescent model.

**Figure 1.**
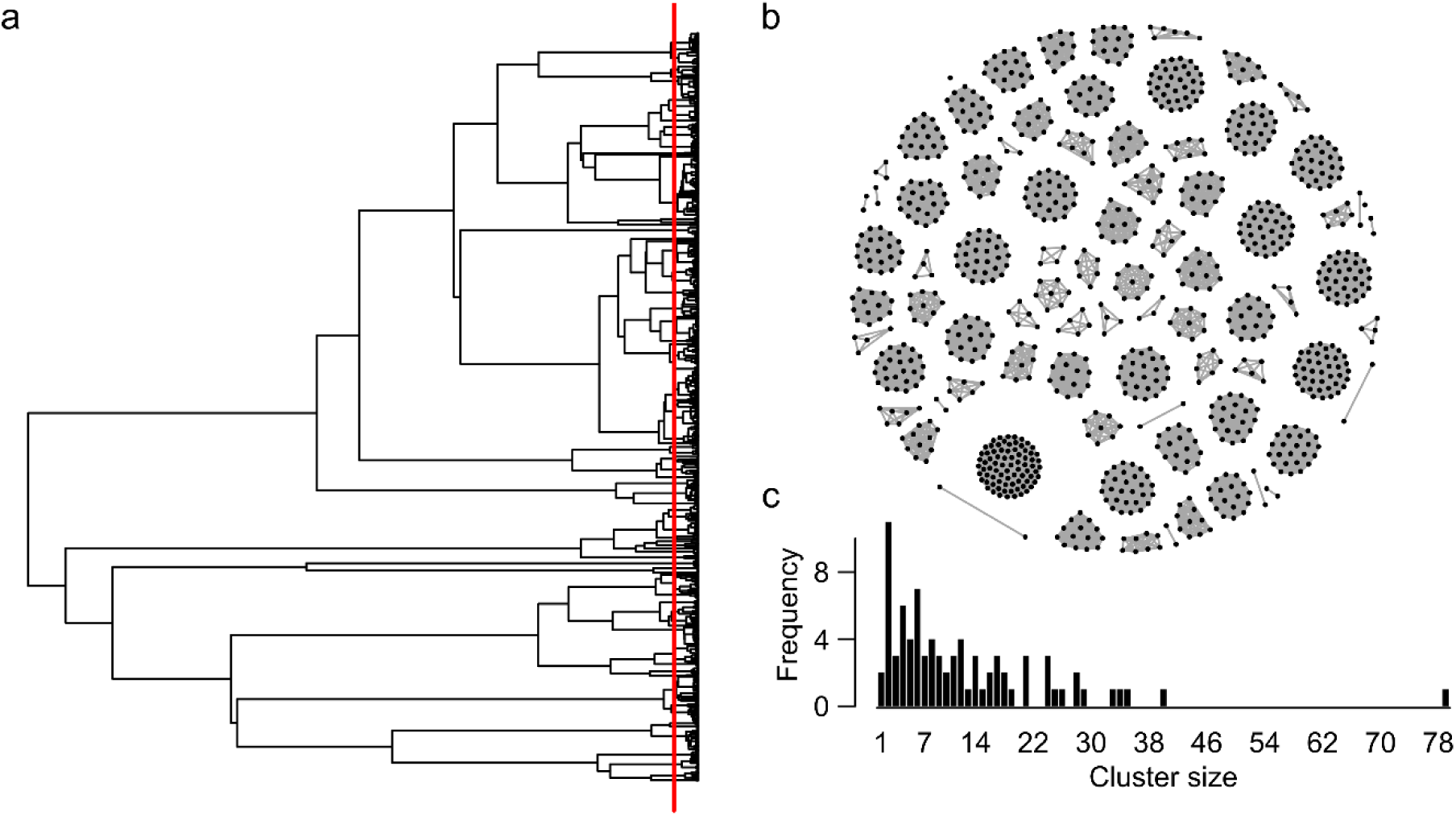
Identifying clusters using a threshold. Clusters from a phylogenetic tree (a) with the threshold cut off marked by the red vertical line, can also be represented by an undirected network (b) or size (component) distribution (c).

The main advantage of clustering sequences from a large database is that it is fast and scalable [6], helping to downsize large sequence databases to groups of particular interest for further in-depth analyses that would be computationally infeasible on the database as a whole. When we have meta-data about clustered individuals, such as potential risk behaviours and clinical results, then clusters can help identify trends between individuals linked within the same cluster and elucidate where to target interventions. Interest also lies in whether these clusters may represent distinct sub-epidemics associated with higher transmission rates. If this were the case, then clusters can be treated separately from the dataset as a whole. However, clusters of related sequences can arise by chance even in the absence of any transmission heterogeneities in the dataset, simply due to the asymmetric way that random trees branch, and can also be driven by oversampling. In this case, treating a cluster of sequences as distinct may be misleading.

Biases can also arise regardless of whether or not clustering is present. Kuhner, Yamato and Felsenstein have previously showed that estimates of the growth rate in coalescent models of exponential growth are biased upwards [15]. Two factors they attributed this to were the nonlinearity of the relationship between the growth rate and the coalescence times, and the censoring of the of the first coalescence event due to sampling. Censoring has also been found to be an issue in non-parametric estimates of the effective population size over time, with false slowing of epidemics detected near the present [16].

In this paper, we investigate the problems of using phylogenetic clusters for phylodynamic inference, motivated by the large national database scenario. Identifying a sub-epidemic with different transmission dynamics relative to the rest of the epidemic early in its development is essential for intervention to have the greatest impact. One way to do this is to consider estimates of epidemiological parameters in transmission clusters to provide insights into the ancestral dynamics [8], and identify whether clusters truly represent sub-epidemics with different transmission patterns. We focus on two main modelling frameworks, the coalescent and birth-death-sampling process, quantifying parameter estimates of the effective population size and growth rate of an epidemic. For the coalescent models, we also implement a model that incorporates censoring for the first coalescence event.

## Methods

### Simulating phylogenies

Coalescent phylogenies were simulated in GENIE v3.0 [17], with the population size in the present, *N(0)* equal to 1000 generations, and the exponential growth rate, ***r***, equal to 0.5. Trees with 200, 400, 600 and 800 tips were simulated under each scenario, with 1000 replicates at each sample size. A further 500 phylogenies were simulated with 500 tips, *N(0)* of 1000 generations and ***r*** equal to 0.01, 0.05, 0.1, 0.5, 1.0 and 5.0 to test the effect of varying the growth.

Birth-death trajectories were simulated using MASTER v 4.1.3 [18], with a birth rate of 0.2 and death rate of 0.1 per unit of time, and an initial population size of one. We again simulated 1000 phylogenies of 200, 400, 600 and 800 tips, with samples taken homochronously after 70 time units. Therefore, in contrast to the coalescent case, more tips represent a higher sampling proportion rather than total population size. We removed any simulations that failed to reach the target sample size.

### Cluster identification

The phylogeny was first converted into a matrix of pairwise distances between tips. Samples were deemed to be in the same cluster if the total branch length, or patristic distance [19], between them was less than the threshold distance. In the example in Figure 1, this is indicated by a red line. The exact threshold depended on the modelling scenario, and was chosen to ensure that the largest cluster used in all further analyses had enough tips to provide useful estimates, but did not include the full tree. Therefore, the star-like trees of the growing populations required thresholds relatively deeper in the phylogeny so as to maintain largest clusters that were not just singletons and pairs. In the constant coalescent simulations we used a threshold of 500 time units, with 15 in the exponential case when ***r =*** 0.5. For the extra exponential phylogenies with 500 tips, thresholds were chosen so as to produce similar distributions for the maximum cluster size (Supplementary Figure 1). In the birth-death scenario, we used a threshold of 50 time units.

To assess the impact of the threshold value, we also looked at two other scenarios. To investigate the effect of conditioning on the largest cluster size at a particular threshold distance, we chose an internal node at random, and all tips linked by that as their most recent common ancestor were deemed to be in the cluster. Internal nodes were resampled if the resulting cluster had less than four tips. This is denoted the ‘random subtree’ method in what follows. We also dropped tips at random throughout the tree (the ‘randomly dropping tips’ method), leaving the same size tree as the largest cluster, to assess any bias in estimates that would be expected from simply having a smaller phylogeny.

### Model fitting and parameter estimation

All models were fitted using maximum likelihood in **R** [20]. For the coalescent simulations, we fitted the constant (***N*(*t*) = *N*(0)**), exponential (***N*(*t*) = *N*(0)e^−rt^**) and logistic (***N*(*t*) = *N*(0)**(**α + (1 — α)e^−rt^)**) demographic coalescent models, where *N(0)* is the initial population size, ***r*** is the exponential growth rate and **α** is the population size at ***t = ∞*** as a proportion of ***N(0)***. Optimization was performed via the BOBYQA algorithm [21] as implemented in the minqa package in **R** [22] using the genieR package available on GitHub (available at: https://github.com/xiangfstats/GenieR). The model with the best fit was decided by taking the model with minimum Akaike Information criterion (AIC) [23].

For the birth-death model, we followed the fitting procedures given in Section 3 of Volz and Frost [24]. The likelihood was maximised using the Nelder-Mead method [25] implemented in the bbmle package in **R** [26]. We assume the death rate is known and fixed at 0.1, and estimate the birth rates and sampling proportion. This makes sense epidemiologically, since these are often parameters of interest during outbreak situations, and independent clinical information can be used for the death rate.

The percentage change in parameter estimate is calculated as:

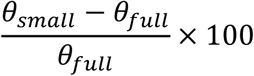

where ***θ_full_*** is the estimate in the full phylogeny, and ***θ_small_*** is the estimate from the downsampled tree (i.e. identified cluster, randomly sampled subtree or from randomly dropping tips). When this value is negative, the smaller tree underestimates the true parameter, and when it is positive, the smaller tree overestimates the full tree parameter.

### Censored Likelihood

To investigate the effect of left censoring of the first coalescent event, we implemented a censored likelihood. For all other coalescence events, we can calculate the exact waiting times from the tree. However, for the first coalescence event, the observed time is cut off by the sampling of the lineages below it. Under the standard coalescent, the probability density function for the time to the next coalescence is:

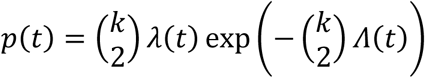

where *k* is the number of lineages, 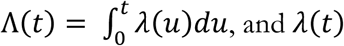 is the rate of coalescence relative to that in the present. Therefore, the probability that the first coalescence event has a time, ***C*_1_** greater than the observed ***t*_1_** is given by:

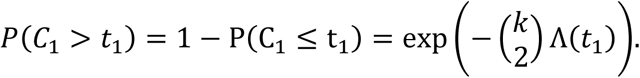

The total likelihood is given by the product of all the probabilities of the coalescence events and sampling events. Thus, the censored likelihood is different from the standard coalescent likelihood by a factor of 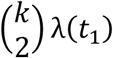.

## Results

### Constant coalescent model

We considered coalescent phylogenies simulated under two demographic scenarios: constant and exponential growth.

In the constant coalescent, the estimated effective population size of the full population was underestimated when considering the false-positive clusters (Figure 2). The effect was similar whether the false-positive cluster was found using a threshold, shown in Figure 2a, or using a random subtree (Figure 2b). The bias was strongest when considering clusters containing the smallest proportion of the full tree, reducing as the cluster size approached that of the full tree. This makes sense, since in a coalescent model, the effective population size is proportional to the branch lengths in the tree. By analysing the cluster without the remaining ancestral context, the depth of the tree has been lost. When trees are down-sampled to the same number of tips at random, then the full depth of the tree is less likely to be lost and the estimate of the full tree more likely to be recovered (Figure 2c). What is perhaps more surprising is the linear nature of the relationship between proportion of tips in the cluster or subtree, and the reduction in estimate of the full effective population size (Table 1). For every 10% (i.e. a proportion of 0.1) more tips present in the cluster relative to the full tree, the accuracy of the effective population size increases by around 9% in the constant case, and this is consistent regardless the number of tips in the tree.

**Figure 2.**
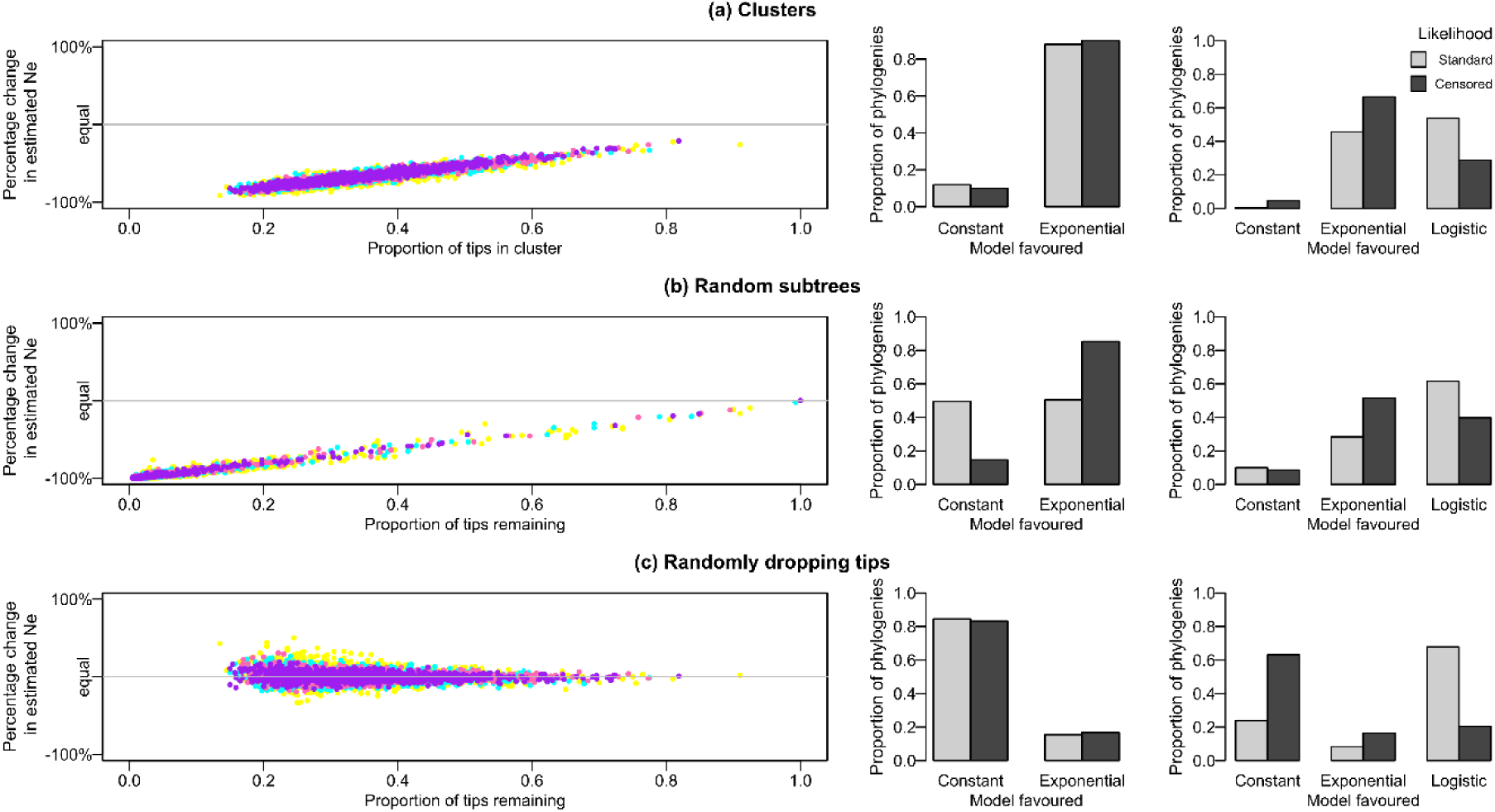
The effect on estimates of the population size in a constant coalescent when considering (a) clusters, (b) random subtrees and (c) tips dropped at random. Colours represent the number of tips in the tree: yellow = 200, green = 400, pink = 600 and purple = 800. The bar plots show the proportion of phylogenies favouring the constant versus exponential (middle column) or constant versus exponential and logistic models (rightmost column) for the standard (light grey) and censored (dark grey) likelihoods.

**Table 1.**
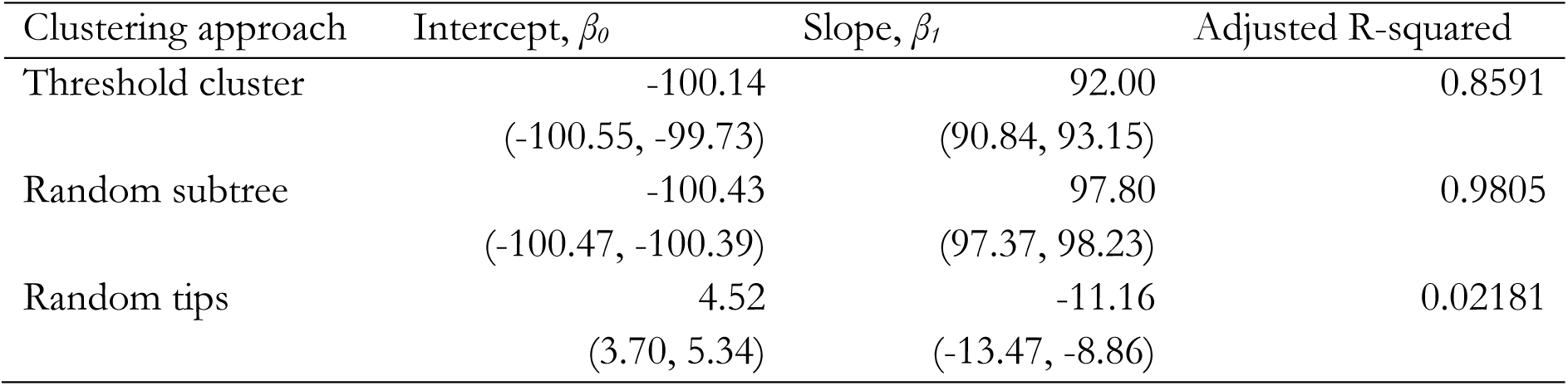
Regression estimates and 95% confidence intervals for the constant coalescent simulations in the form of y = *β_0_* + *β_1_*/x, where y is the percentage change in estimated *Ne*, and x is the proportion of tips in the cluster.

| Clustering approach | Intercept, $\beta_0$ | Slope, $\beta_1$ | Adjusted R-squared |
| --- | --- | --- | --- |
| Threshold cluster | -100.14<br>(-100.55, -99.73) | 92.00<br>(90.84, 93.15) | 0.8591 |
| Random subtree | -100.43<br>(-100.47, -100.39) | 97.80<br>(97.37, 98.23) | 0.9805 |
| Random tips | 4.52<br>(3.70, 5.34) | -11.16<br>(-13.47, -8.86) | 0.02181 |

Although of theoretical interest, the effective population size of a cluster relative to the full tree is less useful in practice. The effective population size is hard to interpret in absolute terms with respect to the number of infected individuals, as it is confounded by the transmission rate [27,28]; however, exponential growth rates as well as overall demographic patterns are well captured when sampling is random. Therefore, we also performed model fitting to the false clusters, to see if they most resembled a constant, exponential or logistic population.

Surprisingly, 67.8% of true constant trees were wrongly fitted as having logistic growth when dropping tips at random, which should have little or no effect on the demographic model. When we instead did model fitting using the censored likelihood, this figure was reduced to 20.4% (constant: 63.2%; exponential: 16.4%). Using clusters biased the model selection further: when comparing between the constant and exponential, the constant model was favoured only 12.0% of the time for clusters compared to 84.6% for trees with randomly dropped tips (censored likelihood: 10.0% vs 83.2%). Adding the logistic model caused the constant model to be favoured for only 1% of the clusters, versus 23.9% of trees with randomly dropped tips (censored likelihood: 4.6% vs 63.2%).

This is particularly problematic when it comes to phylodynamic inference for identifying intervention strategies. Using a metapopulation coalescent [29], it can be shown that exponential growth of the viral population is equivalent to a susceptible-infectious (SI) epidemiological model in the host, and the logistic curve is equivalent to the susceptible-infectious-susceptible model (SIS).

### Exponential Coalescent Model

The results for the effective population size of the exponential model were broadly similar to that of the constant model (Supplementary Figure 2), though with much more noise. The long branch lengths in the present relative to the past, typical of exponential tree shape, meant that the effective population size tended to be overestimated when using particularly small random subtrees, and also when tips were randomly dropped. The constant demographic model was rejected for all cases (Figure 3), with the exponential model rejected for 47.9% of the clusters but only 9.3% of the trees with randomly dropped tips (censored likelihood: clusters 41.3% vs dropped tips 5.0%).

**Figure 3.**
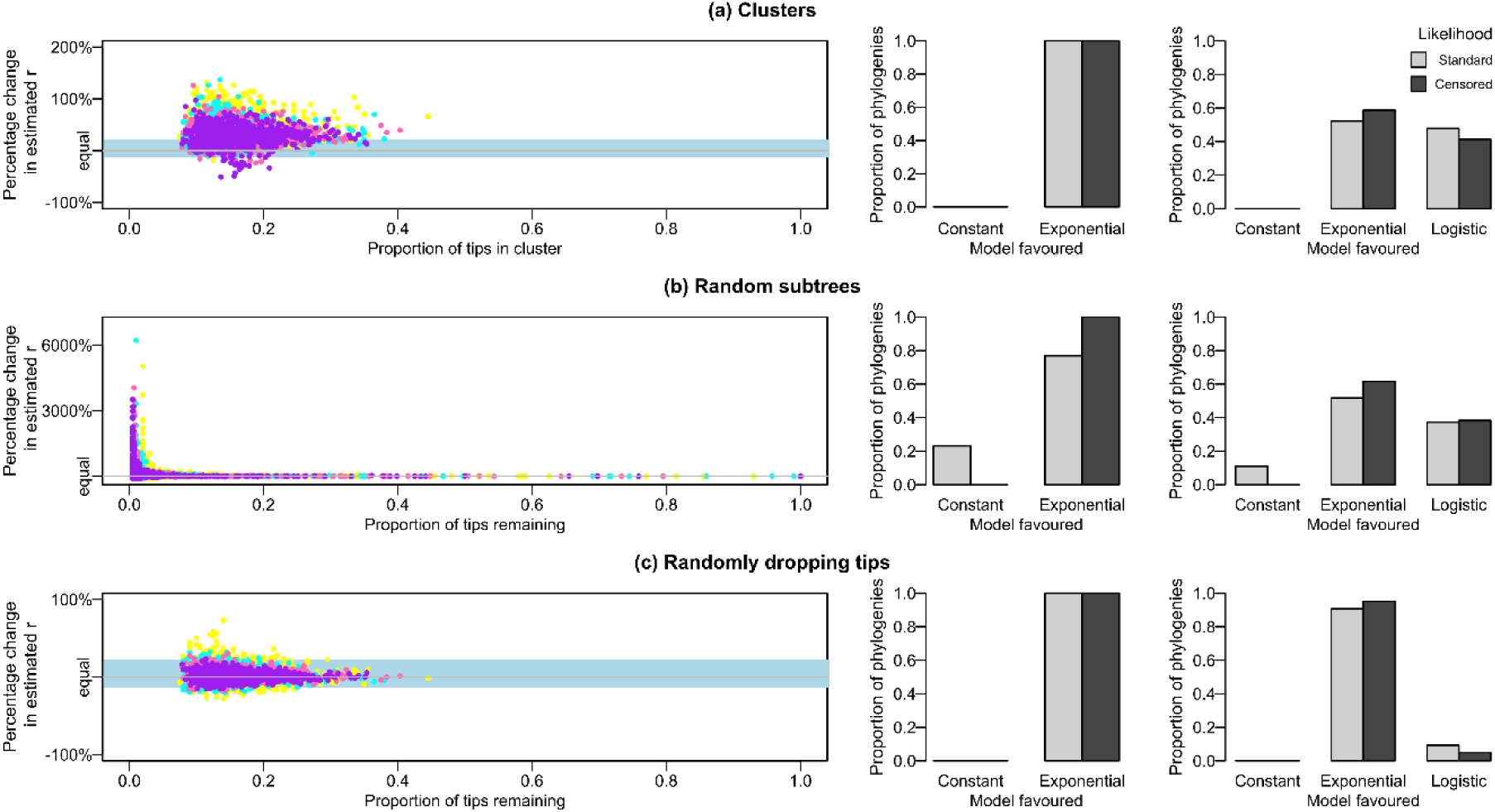
The effect on estimates of the growth parameter, ***r***, in an exponential coalescent when considering (a) clusters, (b) random subtrees and (c) tips dropped at random. Colours represent the number of tips in the tree: yellow = 200, green = 400, pink = 600 and purple = 800. The bar plots show the proportion of phylogenies favouring the constant versus exponential (middle column) or constant versus exponential and logistic models (rightmost column) for the standard (light grey) and censored (dark grey) likelihoods.

For growing populations, using false clusters might seem less problematic than in the simplest constant case if there was other evidence, such as epidemiological data, also suggesting a growing population. However, Figure 3 shows that the estimates of the growth parameter, *r*, tend to be biased upwards in clusters relative to the true value in the tree. This overestimation is beyond that which occurs when tips are dropped at random (Figure 3c). The bias calculated from these randomly dropped tip trees represents the expected bias simply due to having a smaller tree, and has a median value of 1.19%, with 2.5 and 97.5 percentiles of −13.74% and 22.03% respectively. For comparison purposes, these extremes are represented by the light blue panels in Figure 3a and Figure 3b.

When taking clusters in these trees (Figure 3a), the median growth parameter bias is 33.27%, with 94.52% clusters having more extreme bias than the same size tree obtained by dropping tips at random. More generally, points are estimated to fall above light blue box 73.58% percent of the time. A slightly lower figure is obtained for the random subtrees (63.1%), where the smaller subtrees give much more varied estimates overall. The pattern of overestimation is much more pronounced for smaller exponential growth rates, with 99.6% of clusters falling outside of the 2.5–97.5% interval from randomly dropping tips when ***r*** = 0.01, but less so for larger ones, for example only 66.0% of clusters fall outside the equivalent interval for ***r*** = 5 (Figure 4).

**Figure 4.**
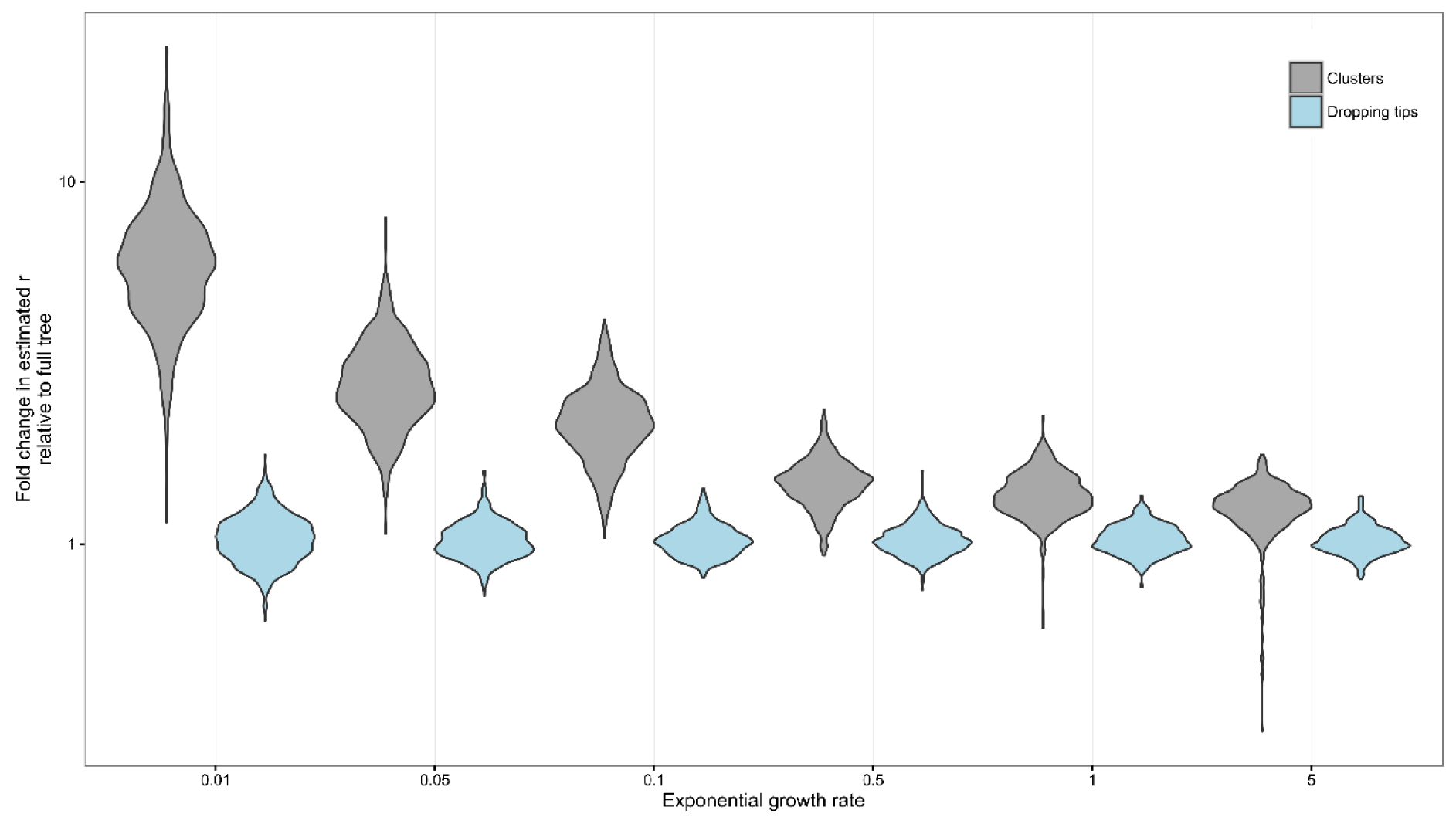
Comparison of fold change in estimated ***r*** for downsampled trees relative the full tree estimate for clusters (grey) and dropping tips (blue). Thresholds were chosen to obtain similar maximum cluster sizes across the different values of ***r*** (shown in Supplementary Figure 1).

### Birth-Death Models

The birth-death results look very similar to those of the exponential coalescent. Like ***r***, the exponential growth parameter, the birth rate, *b*, was overestimated in the clusters (Supplementary Figure 3). When randomly dropping tips, the median overestimate is 1.84%, with the 2.5 and 97.5 percentiles –13.38% and 42.92% respectively (again, represented by the light blue panels in Supplementary Figure 3). The median overestimate rises to 22% (-23.33, 457.90) for the randomly chosen clades, and 31.5% (0.69, 94.22) using threshold clustering. Making a direct comparison between the dropped tips and cluster results, since they have the same down sampled size, the cluster result bias is more extreme (whether over or underestimating) than of the dropped tip tree in 90.8% of the phylogenies.

The population size, calculated using the size of the tree and the estimated sampling proportion, is also underestimated (Supplementary Figure 4). The line on the graphs shows where *y*=*x*, and when dropping tips at random, the full tree estimate is recovered. Clusters and random subtrees underestimate the majority of the time, with results generally noisier in the latter: the cluster overestimates the population for 1.4% of phylogenies, versus 5% for random clades and 57.95% when dropping tips at random. Similar results were obtained when using the tree likelihood conditioned on the number of tips in the phylogeny (as given by Equation 3 in [30]).

## Discussion

Early identification of new epidemics is essential for interventions to be most effective. This is even more important in a highly clustered epidemic, where, unless interventions are targeted appropriately, the infection will continue to spread regardless of its underlying transmissibility [12]. In this paper, we show that whilst clusters can be useful for finding affinities between closely related samples, there are some caveats when using them to understand demographics in subpopulations. Most importantly, false clusters can masquerade as growing epidemics.

Clusters have previously been used in the literature to downsample large datasets and look at the fit of demographic models [13,31,32]. For example, Hué et al. [13] identified six clusters of UK origin, and found all of them to be best fitted by a logistic growth model with an doubling time of approximately a year during the initial exponential growth stage. Our study shows that it is feasible that these clusters are not growing at all, and are simply an artefact of using a threshold to downsample a large database taken from a randomly mixing population.

Identifying clusters is not the only problem when inferring population dynamics, however. Previously noted biases were reiterated when looking at our trees downsampled by randomly dropping tips: the over-estimation of the exponential growth rate parameter and the false fitting of the logistic model over the constant [15,16]. De Silva et al. [16] showed that there exist biases in non-parametric estimates of the population size, with false slowing of the exponential growth parameter near to the present. In their paper, they suggested that this is due to left censoring backwards in time and the relative time between the first coalescence and sampling events. Here, the majority of true constant downsampled trees were wrongly fitted as having logistic growth when using standard parametric models. We implemented a censored likelihood for the first coalescent event which went some way to improving the model selection, suggesting that demographic biases from censoring are not limited to non-parametric methods (c.f. the conclusions of de Silva et al.). The censoring appeared to have less effect on the model selection in clusters, probably due to the overall reduced tree height.

Clustering itself has a number of disadvantages, particularly pertaining to the arbitrary choice of threshold. Most clustering methods are defined by non-parametric methods, that is, not by a model, and neglect the rest of the information deeper in the phylogeny’s ancestry, with small thresholds biasing cluster membership to recent infection events. Non-epidemiological factors such as sampling are also an issue [33]; oversampling in an area or time period can skew results, and unsampled individuals mean that a point source transmission to a single cluster cannot be excluded.

When clusters truly represent different subpopulations, it makes sense computationally to analyse them outside the background of the rest of the database. One obvious problem is that large databases will not have one single background demographic model as considered here. Instead, there may be one or more clusters of varying age, demography and viral viability. Effect on demography estimates aside, the phylogenetic clustering is ill-equipped for dealing with these. A recent simulation study by Poon [34] showed that many clustering methods cannot detect heterogeneity in transmission rates between subpopulations. Instead, they focus on individuals with a short waiting period between transmission and diagnosis. This has further ramifications on targeting interventions, as clusters may well be identifying subpopulations that are already seeking medical attention.

Methods for identifying up-and-coming epidemics from within a large database need to be able to look for growth above and beyond the biases we have shown exist. Clustering approaches have the advantage that they are reasonably fast even on relatively large datasets [6], especially compared to alternative approaches such as the structured coalescent. There have been a number of recent developments which have made the structured coalescent more amenable for this sort of analysis, however fitting many demographic models to unknown stratifications in the sample remains difficult [35–37]. One way to understand the population background in the tree for comparison to an identified cluster would be to use information in the clades outside of the cluster to form a null distribution. The clustering method of Prosperi et al. [7] uses genetic distance between tips for a similar purpose. For phylodynamic inference, a permutation approach such as that in [38] could be used to assess whether observed clustering is due hidden heterogeneity. In addition to asymmetry metrics for measuring heterogeneity across the phylogeny, the clade size distribution and exponential growth or birth rate calculated from different time points (e.g. for the subtree below each internal node) could add additional information.

Until a suitable null distribution or parametric method for identifying clusters is developed, their interpretation will remain problematic for phylodynamics. In particular, we should be cautious when targeting intervention strategies on the basis of clustering, to ensure resources are put where they will have most impact, and not to populations already active in seeking medical help.

## Acknowledgements

This work was supported by a Medical Research Council Methodology Research Programme grant to S.D.W.F (grant number MR/J013862/1). FX is supported by a BBSRC Strategic LoLa grant (grant number BB/L001330/1).

## Supplementary Information

**Supplementary Figure 1.**
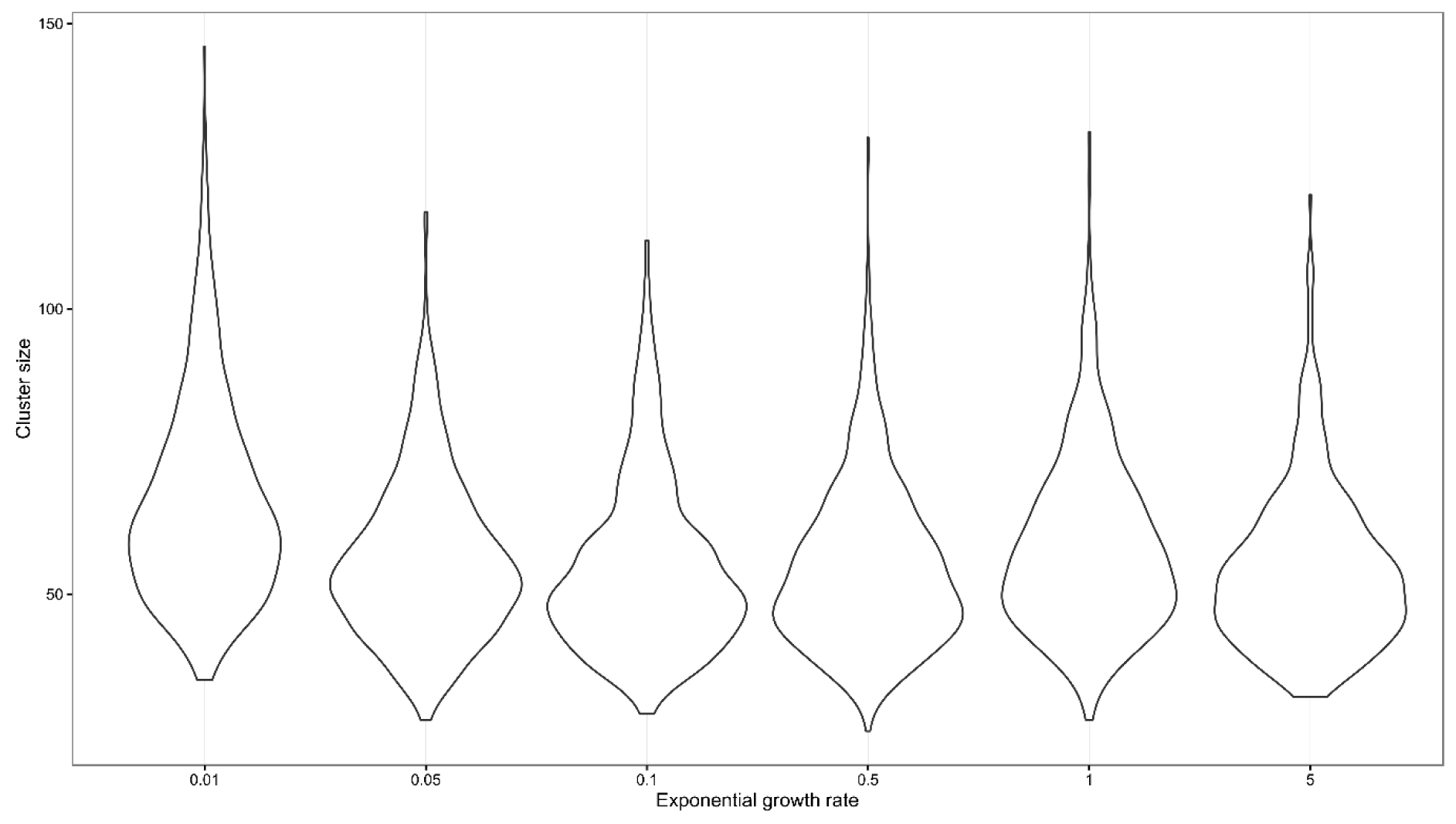
Violin plots showing the maximum cluster sizes used when comparing between different exponential growth rates.

**Supplementary Figure 2.**
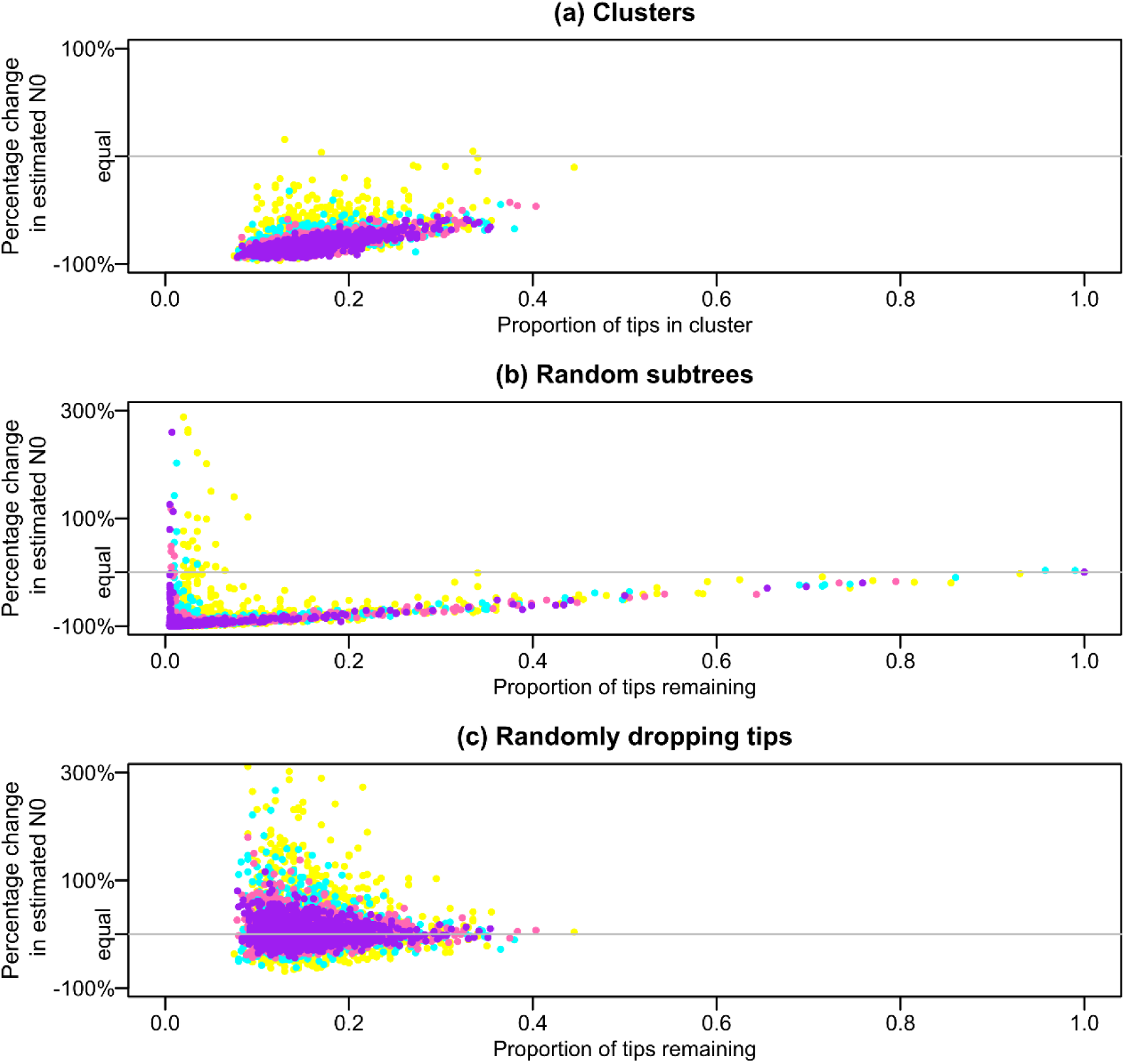
The effect on estimates of the present population size, *N_0_*, in an exponential coalescent when considering (a) clusters, (b) random subtrees and (c) tips dropped at random.

**Supplementary Figure 3.**
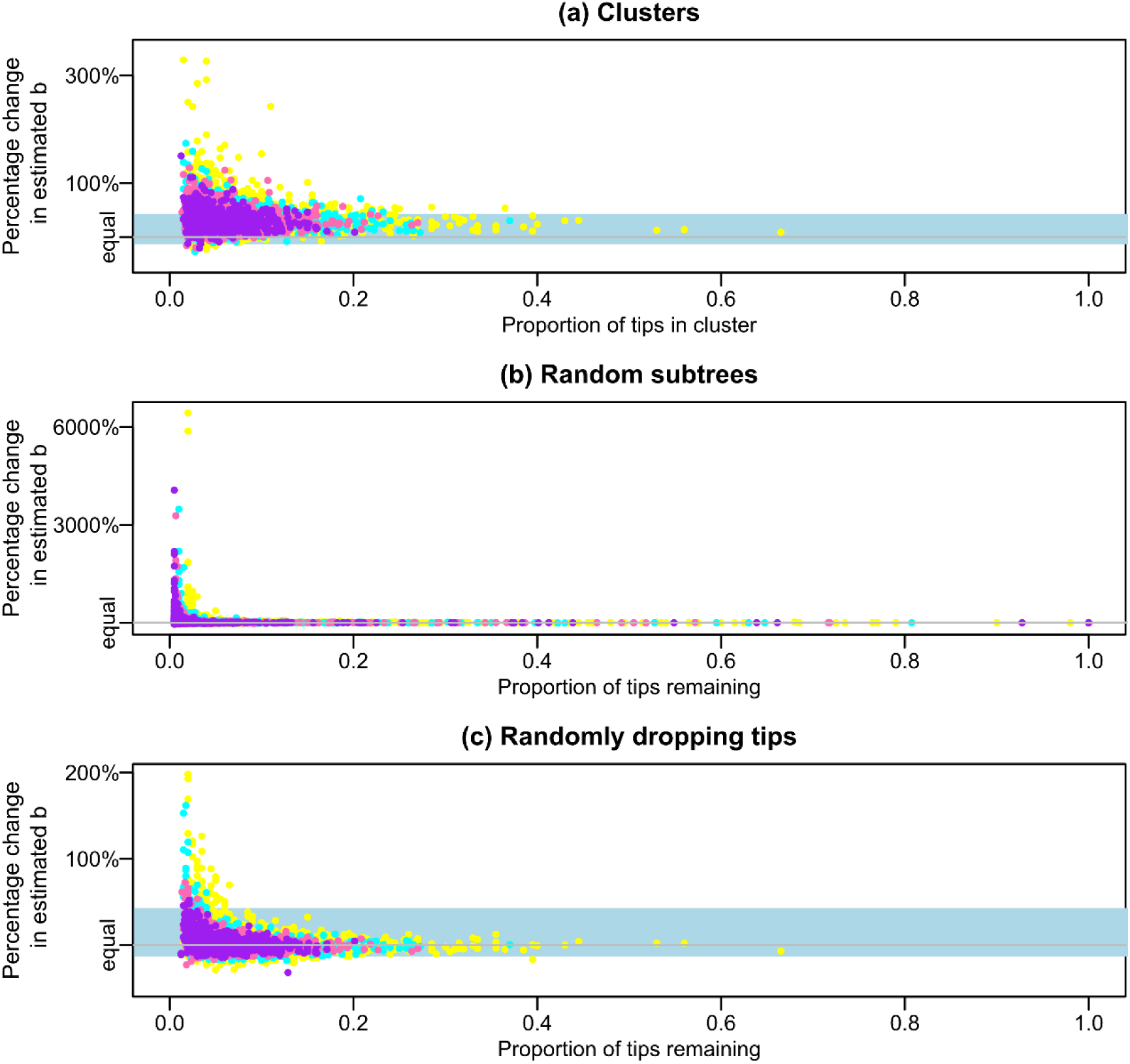
The effect on estimates of the birth rate, *b*, from the birth-death model when considering (a) clusters, (b) random subtrees and (c) tips dropped at random. Colours represent the number of tips in the tree: yellow = 200, green = 400, pink = 600 and purple = 800.

**Supplementary Figure 4.**
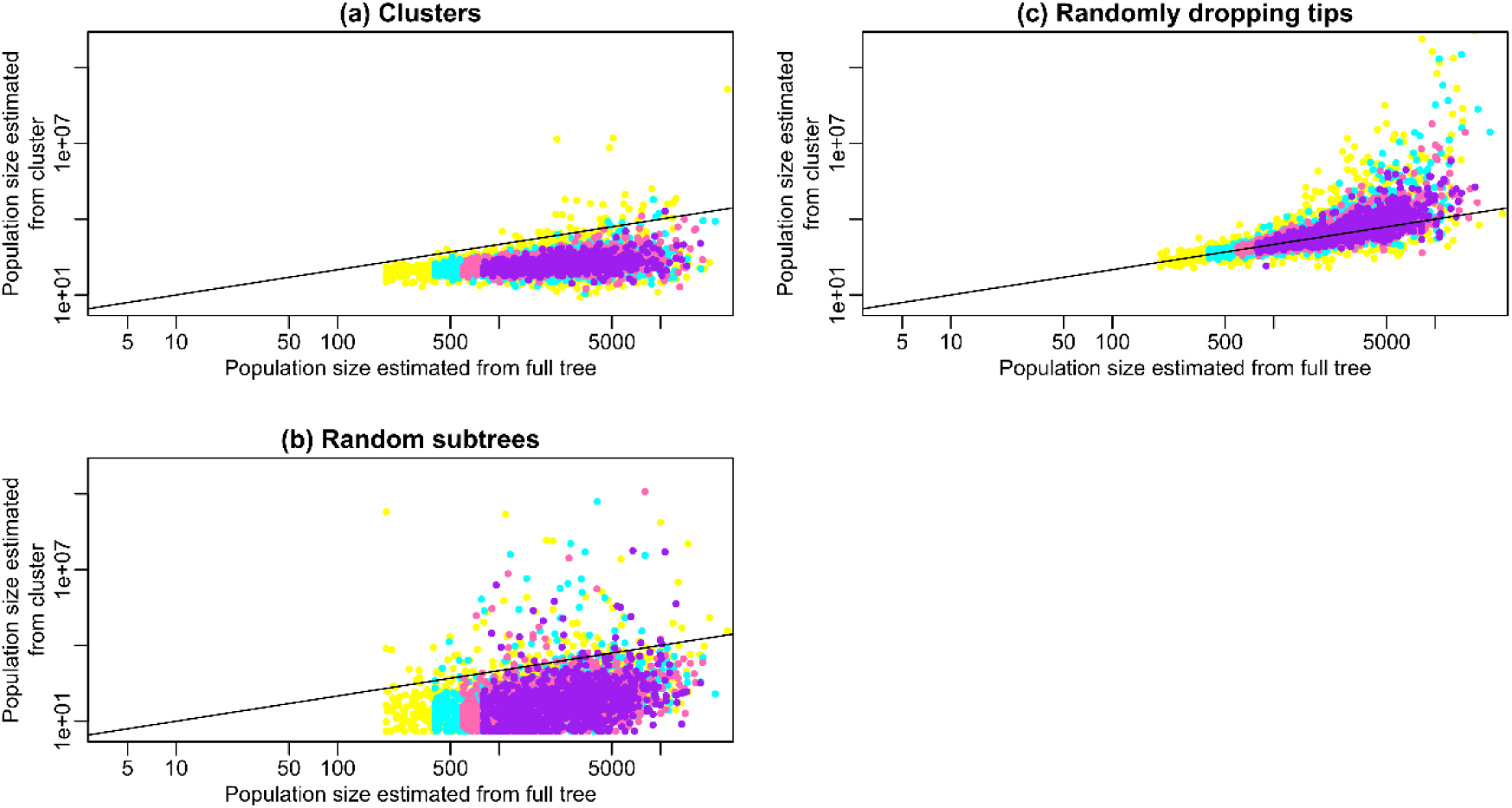
The effect on estimates of the population size as estimated from the sampling proportion and tree size in from the birth-death model when considering (a) clusters, (b) random subtrees and (c) tips dropped at random. The line shows where *y* = *x*. Colours represent the number of tips in the tree: yellow = 200, green = 400, pink = 600 and purple = 800.

